# Increased frequency of CD4^+^ and CD8^+^ follicular helper T cells in human lymph node biopsies during the earliest stages of rheumatoid arthritis

**DOI:** 10.1101/2021.11.10.467883

**Authors:** Dornatien C. Anang, Tamara H. Ramwadhdoebe, Janine Hahnlein, Bo van Kuijk, N. Smits, Krijn P. van Lienden, Mario Maas, Danielle M. Gerlag, Paul P. Tak, Niek de Vries, Lisa G. van Baarsen

## Abstract

**Objectives:** Follicular helper T cells (Tfh cells) provide key B cell help, and are essential in germinal center (GC) formation and (auto) antibody generation. To gain more insight into their role during the earliest phase of rheumatoid arthritis (RA) we analyzed their frequencies, phenotype and cytokine profile in peripheral blood and lymphoid tissues.

**Methods:** Using flow cytometry, we studied the frequency of Tfh and B cells in peripheral blood and lymph node (LN) needle biopsies. Three donor groups were included and compared: healthy controls (HCs), autoantibody positive individuals at risk for developing RA (RA-risk individuals), and early RA patients. *Ex vivo* stimulation of lymphocytes with PMA/ionomycin was performed to assess cytokine secretion by Tfh cells.

**Results:** In blood, the frequency of circular Tfh cells (cTfh) did not differ between study groups. In lymphoid tissue, the frequency of Tfh cells correlated strongly with the frequency of CD19^+^ B cells. Compared to healthy controls, LN samples of RA patients and RA-risk individuals showed more CD19^+^ B cells and more CD4^+^CXCR5^+^ and CD8^+^CXCR5^+^ Tfh cells. These Tfh cells from LNs expressed less IL-21 upon *ex vivo* stimulation.

**Conclusion:** LN tissue of early RA patients as well as part of RA-risk individuals exhibit increased frequencies of Tfh cells correlating with increased numbers of B cells. Interestingly, IL-21 production is already aberrant in the very early at risk phase of the disease. This suggests that Tfh cells may present a novel rationale for therapeutic targeting during the preclinical stage of the disease to prevent further disease progression.

## INTRODUCTION

The presence of autoantibodies years before the presence of clinical signs and symptoms in rheumatoid arthritis (RA) suggest an increase in B cell differentiation towards antibody-producing plasma cells already very early in the disease (1). Such proliferation and differentiation of B cells is supported by CD4^+^ follicular T helper cells (Tfh cells) (2). These Tfh cells derive from naïve CD4^+^ T cells. Following TCR stimulation under the influence of receptor-ligand interactions and cytokine milieu, T cells may upregulate CXCR5, causing them to migrate to the CXCL13 expressing follicular border where they interact with B-cells and receive signals that will potentially drive them into Tfh cells (3). Producing cytokines like IL-21 and IL-4, Tfh cells interact with B cells which results in B-cell differentiation towards short-lived plasmablasts or in B-cell migration into the follicles to contribute to the formation of germinal centers (GC) (4). Migration of Tfh cells into the B-cell follicles further supports GC formation and drives B-cell differentiation into memory B-cells or long-lived antibody-secreting plasma cells. Overall, the net effect of this process depends on the balance between inflammatory and regulatory signals and is tightly regulated to prevent aberrant (auto)immune activation (5).

One of the studies showing the contribution of CD4^+^ Tfh cells in autoimmunity used the sanroque mouse model. Sanroque mice lack a repressor of ICOS, Roquin, resulting in excessive Tfh cell formation and subsequent GC formation. These mice have increased titres of autoantibodies and lupus-like symptoms (6). In human studies, SLE patients also show increased frequencies of blood Tfh cells compared with healthy controls (HCs) which correlate with an increase in circulating autoantibodies (7-10). Similarly, increased frequencies of peripheral blood Tfh cells have been detected in type I diabetes and RA patients (11-15). Since Tfh cells can leave the lymphoid tissues and relocate through peripheral blood they may contribute to B-cell differentiation and formation of tertiary lymphoid structures at sites of inflammation.

While the role of CD4^+^ Tfh cells in autoimmunity is widely accepted, current data regarding a potential role for CD8^+^CXCR5^+^ follicular-like T cells are less clear, and actually point to a dual role. Like CD4^+^ Tfh cells, CD8^+^ Tfh cells express CXCR5, BCL6, IL-21, ICOS as well as PD-1(16-18). An ex vivo study reported the ability of CD8^+^ CXCR5^+^ T cells to induce the apoptosis of CD4^+^ Tfh cells, resulting in inhibition of IL-21 production (19). However, an antibody-enhancing function of these cells has also been reported in virus-infected mice where IL-21 producing CXCR5^+^ICOSL^+^CD8^+^ T cells were shown to enhance the production and class-switching of IgG antibodies, revealing a major role of CD8^+^ Tfh cells in the immune response (20).

Studies into the role of T cells in the pathogenesis of RA in humans have mainly focused on the well-known Th1, Th2, Th17, and Treg subsets (21-23), during established disease, and in cell populations from peripheral blood and inflamed joints (22, 24). Studies of Tfh cells focused on peripheral blood samples or inflamed tissue (4, 25-27), while studies investigating Tfh cells in lymphoid organs during the earliest phases of autoimmunity are lacking. To the best of our knowledge, no study has analysed CD8^+^ Tfh during the various stages of RA. To gain more insight into the initial activation of Tfh cells in lymphoid tissue and in the role of lymph node Tfh cells during the earliest phases of RA, more research is needed.

In this study, we hypothesized that Tfh cells contribute to autoantibody production in the earliest preclinical phases of RA by driving B cell differentiation in secondary lymphoid organs. Using samples acquired by core-needle biopsies of inguinal LNs (28, 29), we analysed and compared the frequencies of CD4^+^ and CD8^+^ Tfh cells in the blood and lymphoid tissue from healthy controls (HCs), autoantibody-positive individuals at risk for developing RA (RA-risk individuals) and early RA patients (30).

## MATERIALS AND METHODS

### Study subjects

Twenty-four individuals at risk for developing RA (RA-risk) were selected. RA-risk status was defined by the presence of IgM rheumatoid factor (IgM-RF) and/or anti-citrullinated protein antibodies (ACPA) positive) in subjects without any evidence of arthritis (30). The median follow-up time was 26.8 months (14.3-39.2 (interquartile range, IQR)) and none of these RA-risk individuals had developed arthritis as yet. We also included 16 early RA patients, based on American College of Rheumatology and European League Against Rheumatism (ACR/EULAR) 2010 criteria (31), who were naïve for disease-modifying antirheumatic drugs (DMARD) and biologicals with a disease duration (defined by having arthritis in any joint) less than 1 year. For comparison, 17 seronegative healthy controls (HCs) were included in the study. The study was performed according to the principles of the Declaration of Helsinki (32), approved by the institutional review board of the Academic Medical Centre of the University of Amsterdam, and all study subjects gave written informed consent. Demographics of all study subjects are listed in table 1.

**Table 1:**
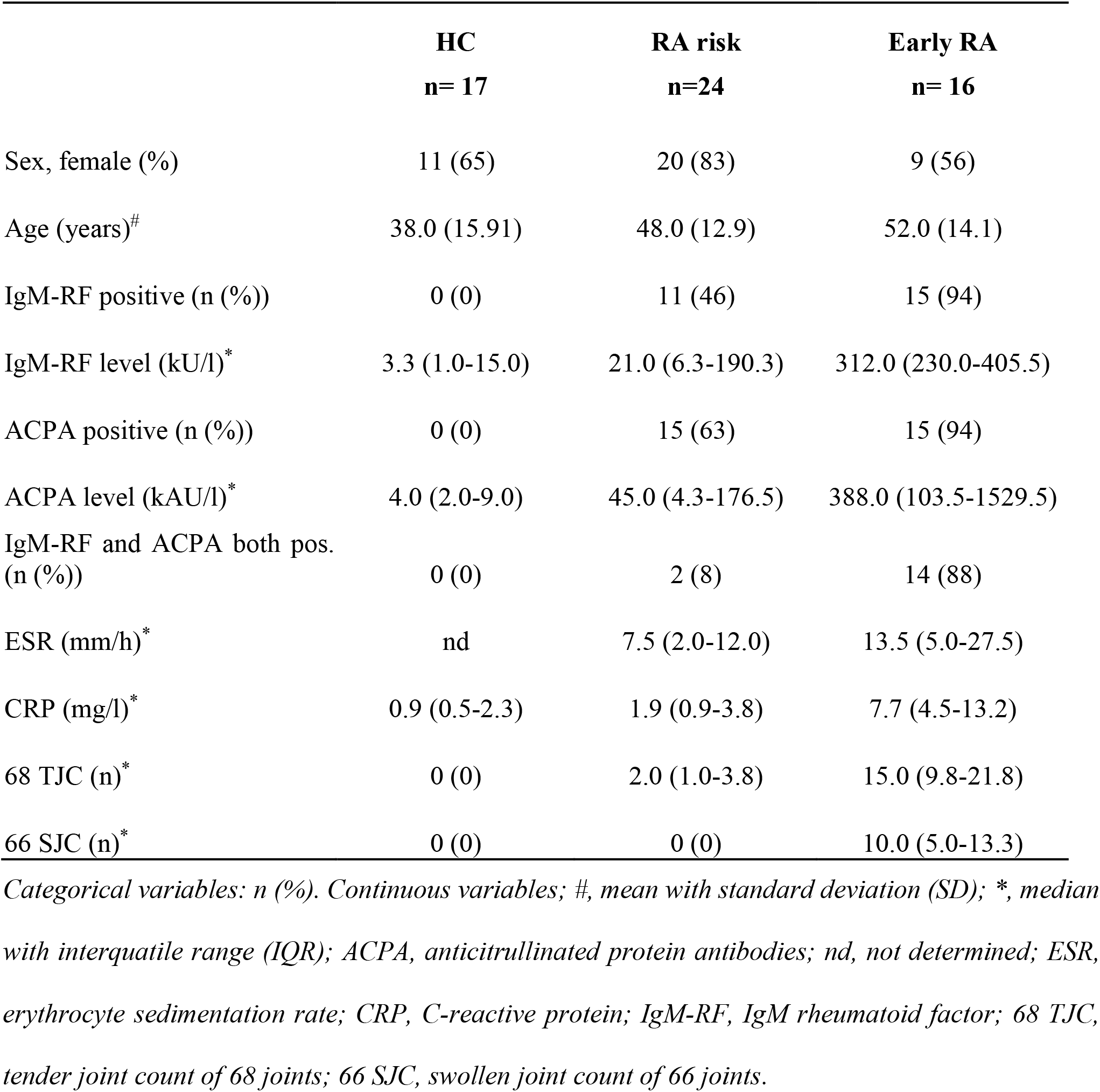
Baseline characteristics of healthy controls (HCs), RA-risk individuals (RA risk) and early RA patients (Early RA)

### Sample processing and cell culture

Ultrasound-guided inguinal LN biopsies were taken and immediately processed for flow cytometry analysis, snap-frozen en bloc in Tissue-Tek OCT compound (Miles, Elkhart, IN) for immunohistochemistry analysis or snap frozen for RNA isolation as described previously (28). For flow cytometry analyses, LN biopsies were put through a 70 μm cell strainer (BD Falcon, San Jose, CA) to obtain a single cell suspension. Peripheral blood mononuclear cells (PBMC) were isolated using standard density gradient centrifugation using lymphoprep (Nycomed AS, Oslo, Norway) and stored in liquid nitrogen until further use. Freshly isolated LN cells and thawed PBMC were incubated in RPMI culture medium (Life Technologies, Thermo Fisher Scientific Inc., Waltham, MA) for 4 hours in the presence or absence of Phorbol Myristate Acetate (PMA) and Ionomycine with Brefeldin A (all from Sigma Aldrich, St Louis, MO) and Golgi stop (BD Biosciences, San Jose, CA). After 4 hours, cells were washed and analyzed by flow cytometry.

### Antibodies and flow cytometry analysis

Cells were stained for 30 minutes at 4°C in PBS containing 0.01% NaN3 and 0.5% BSA (Sigma Aldrich) with directly labelled antibodies against: CXCR5 alexa fluor 488 (clone RF8B2), CCR7 PE-Cy7 (clone 3D12), CD4 APC-H7 (clone SK3), CD8 V450 (RPA-T8), CD3 V500 (UCHT1) (all from BD Biosciences, San Jose, CA), PD-1 PE (J105), CD45 RA (L307.4) efluor450 (all from eBioscience Inc., San Diego, CA). For intracellular cytokine staining we used IL-10 Pe-Cy7 (clone JES3-9D7) (Biolegend, San Diego, CA) and IL-21 alexa fluor 647 (clone 3A3-N2.1) (BD Biosciences). Cells were analyzed on a FACS Canto II (BD Biosciences) and data were analyzed using FlowJo software (FlowJo, Ashland, OR). Data were plotted as frequency of positive cells.

### Statistical analysis

After testing for normality with D’Agostino and Pearson omnibus test, data are presented as mean with standard deviation or median with IQR. Differences between groups were analysed using the Kruskall Wallis test or one-way analysis of variances (ANOVA). Correlations were calculated with Spearman’s Rank Correlation Coefficient. All statistical analyses were performed using GraphPad Prism Software (version 6, GraphPad Software, Inc. La Jolla, CA). P-values <0.05 were considered statistically significant.

## RESULTS

### The frequency of peripheral blood CD4^+^ and CD8^+^ circulating follicular helper T cells is comparable between healthy controls, RA-risk individuals and early RA patients

To analyse the frequency of peripheral blood CD4^+^ and CD8^+^ cTfh cells in HCs, RA-risk individuals and early RA patients we first analysed the total number of CXCR5^+^ and PD-1^+^ cells within the CD4^+^ and CD8^+^ T cells (see gating Figure 1A and supplementary figure 1 (negative control)). The frequencies of CD4^+^CXCR5^+^ and CD4^+^PD-1^+^ T cells as well as CD8^+^CXCR5^+^ and CD8^+^PD-1^+^ T cells were on average comparable between the three study groups (Figures 1B and C). Within the CD4^+^CXCR5^+^ T cell subset, the CCR7^low^PD1^high^ cells have been described as active Tfh cells and the CCR7^high^PD1^low^ as quiescent Tfh cells (14). In our analysis the frequency of active (CCR7^low^PD-1^high^) and quiescent (CCR7^high^PD-1^lo^) cTfh cells within the blood CD4^+^CXCR5^+^ and CD8^+^CXCR5^+^ cells was comparable between the three study groups (Figures 1D and E). No significant correlations between the frequencies of various cTfh cells were present with age or autoantibodies detected in blood (data not shown).

**Figure 1.**
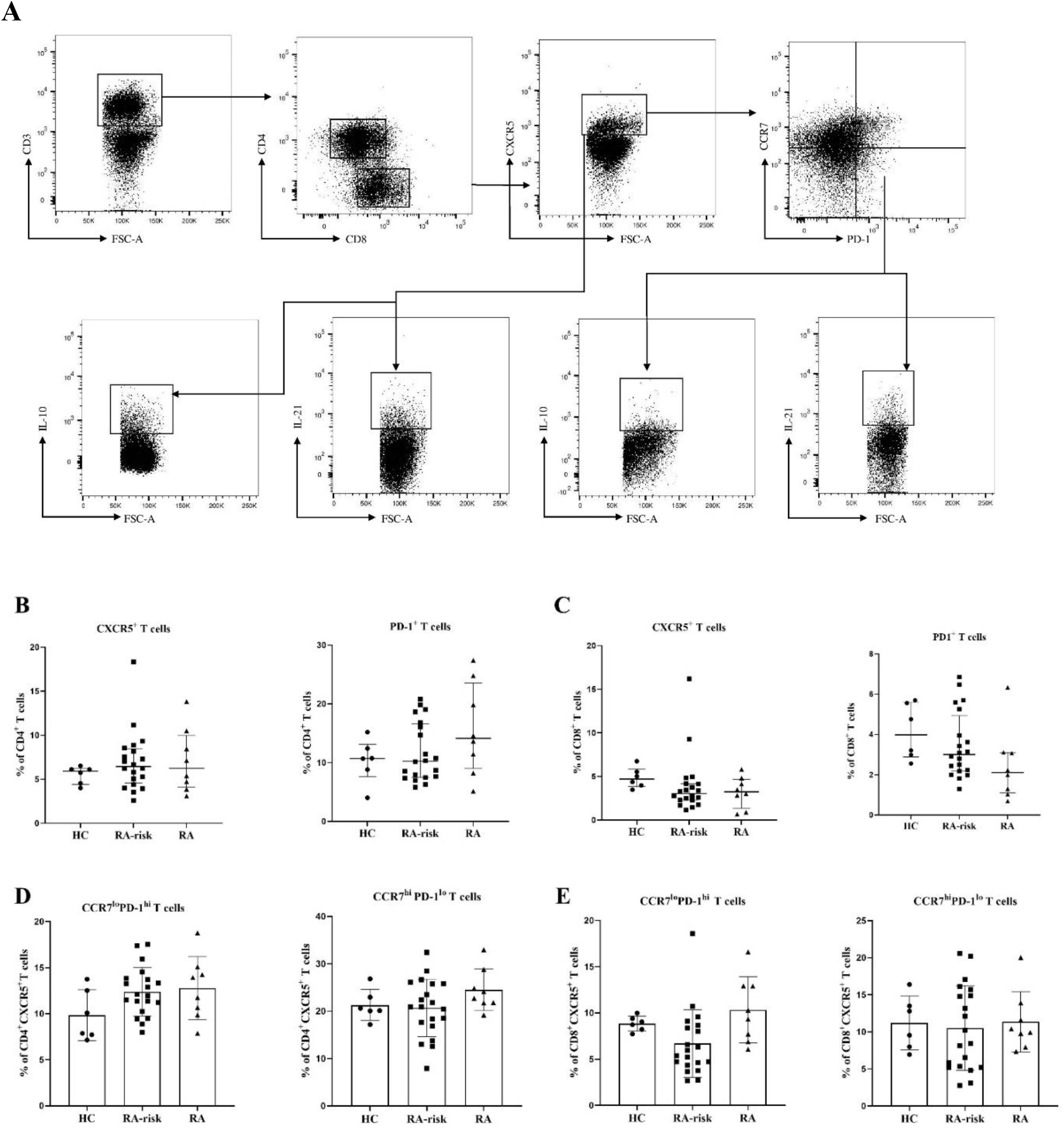
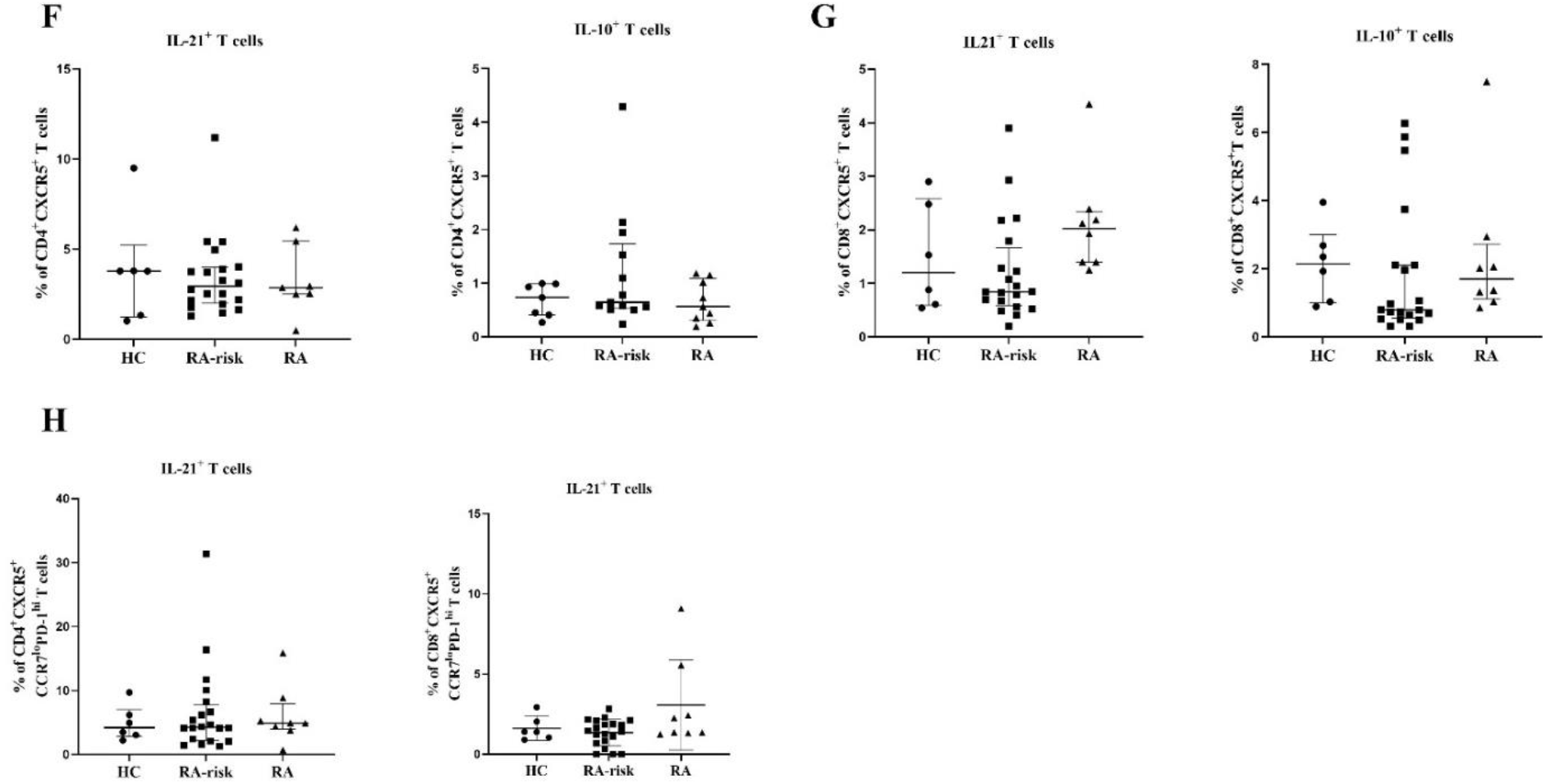
Analysis of circulating follicular helper T cells in peripheral blood samples. **(A)** Gating strategy for cTfh cells in PBMCs using markers for CD3, CD4, CD8, CXCR5, CCR7, PD-1, IL-21 and IL-10. After gating for single cells, CD4^+^ and CD8^+^ T cells were gated within the CD3^+^ population and further characterized. **(B, C)** The frequencies of CXCR5^+^ and PD-1^+^ cells within CD4^+^ and CD8^+^ T cells were analysed in PBMC samples collected from healthy controls (HC n=6), RA-risk individuals (RA-risk n=20) and early RA patients (RA n=8). **(D, E)** the frequencies of blood CCR7^low^ PD1^high^ cTfh and CCR7^low^ PD1^high^ cTfh within the CD4^+^ CXCR5^+^ and CD8^+^ CXCR5^+^ populations are plotted. **(F, G, H)** the frequency of IL-21 and IL-10 producing cells within the indicated T-cell subsets are shown. Not normally distributed data are presented as median with IQR and, normally distributed data are presented in box plots with error bars representing mean with SD. For statistical analysis Kruskall Wallis or one-way ANOVA (when appropriate) was performed. Significant differences were indicated as *p<0.05 or **p<0.01. All symbols represent data from single individuals (• healthy controls (HC), ^▄^ RA-risk individuals (RA-risk), Δ early RA patients (RA).

Next we analyzed the capacity for cytokine production in blood cTfh cells upon ex vivo stimulation with PMA/ionomycin. The frequency of CD4^+^CXCR5^+^IL-21^+^, CD4^+^CXCR5^+^Il-10^+^, CD8^+^CXCR5^+^IL-21^+^ and CD8^+^CXCR5^+^IL-10^+^ T cells was on average comparable between the three study groups (Figures 1F and G). As expected, the frequency of peripheral blood active-Tfh cells producing Il-21 is higher than the frequency of quiescent-Tfh cells. Finally, the frequency of IL-21^+^ cells among active CD4^+^CXCR5^+^CCR7^low^PD-1^high^ and CD8^+^CXCR5^+^CCR7^low^PD-1^high^ was on average comparable between the three study groups (Figure 1H).

Taken together, the frequency of blood CD4^+^ and CD8^+^ cTfh cells is highly variable but on average not different between RA-risk individuals and early RA patients compared with healthy controls (HCs).

### CD4^+^ and CD8^+^ follicular helper T cells are increased in lymphoid tissue of RA patients

Next, we analysed the frequencies of CD4^+^ and CD8^+^ Tfh cells based on CXCR5 expression in LN biopsies (see gating Figure 2A). CXCR5 expression defines LN T cells that are capable of moving towards the follicular border where they can interact with B cells (2). Among CD3^+^ T cells, the frequency of total CD4^+^ T cells was not significantly higher in RA-risk individuals compared to HCs, but significantly increased in early RA individuals compared to HCs (Figure 2B). Among CD4^+^ T cells the frequency of CXCR5^+^ Tfh cells is increased in early RA patients compared with HCs (Figure 2B; p<0.05), while frequencies in RA at-risk individuals are in between. We next evaluated the frequencies of CD8^+^ T cells. Among CD3^+^ cells, the frequency of CD8^+^ T cells was similar between RA-risk individuals and HCs, while it was significantly lower in early RA patients compared to HCs (Figure 2C). Among CD8^+^ T cells, the frequency of CD8^+^CXCR5^+^ Tfh cells was markedly increased (p<0.05) in RA-risk individuals as well as early RA patients when compared to HCs (Figure 2C).

**Figure 2.**
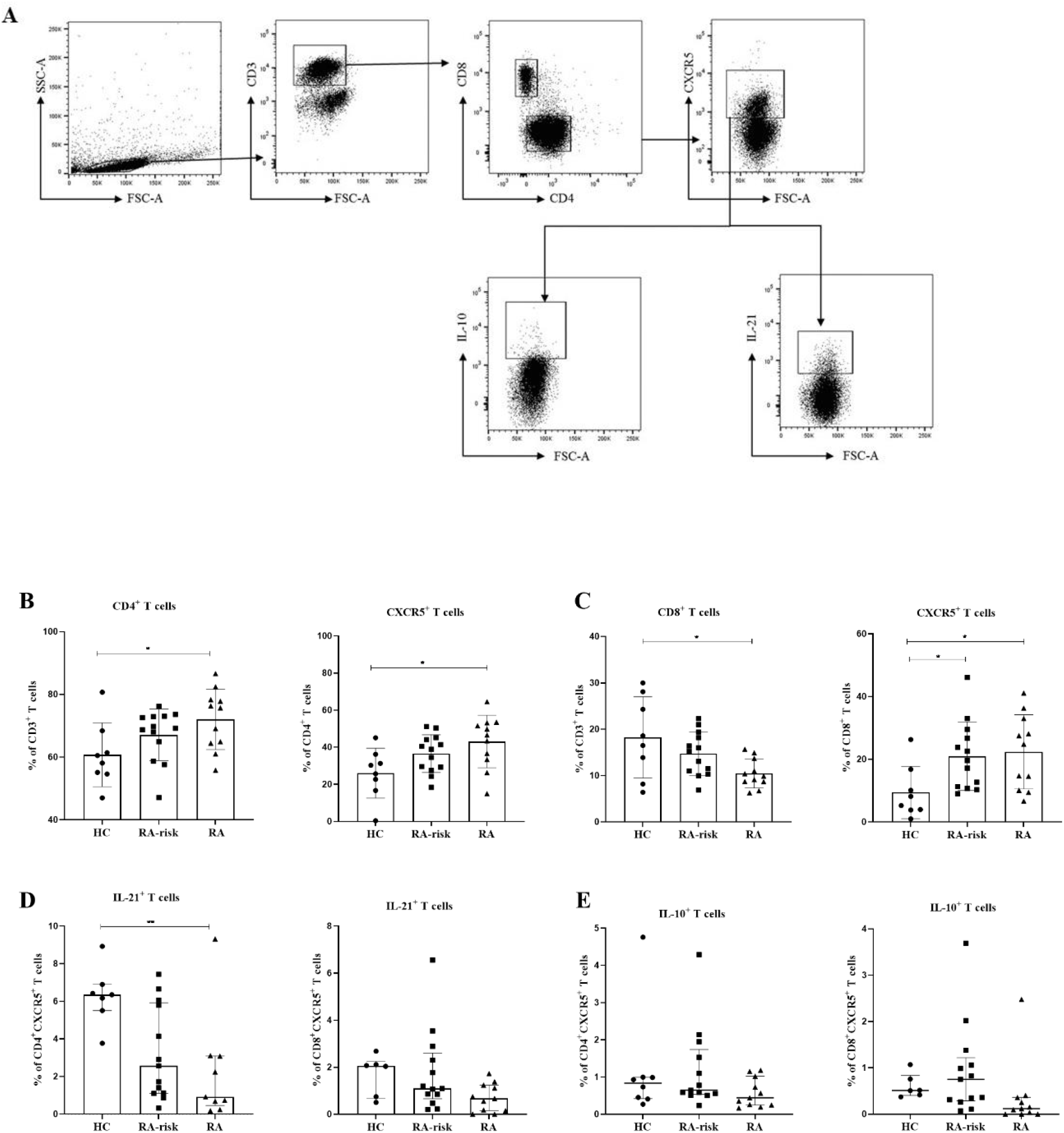
Analysis of Follicular helper T cells in lymph node biopsies. **(A)** Gating strategy for follicular-like T cells in lymph node biopsies using markers for CD3, CD4, CD8, CXCR5, IL21 and IL-10. **(B, C)** Frequencies of CD4^+^, CD4^+^CXCR5^+^, CD8^+^ and CD8^+^CXCR5^+^ T cells in lymph nodes are shown for healthy controls (HC, n=7), RA-risk individuals (RA-risk n=13) and early RA patients (RA n=9). **(D, E)** The frequency of IL-21^+^ and IL-10^+^cells within the CD4^+^CXCR5^+^ and CD8^+^CXCR5^+^ populations are plotted. Not normally distributed data are presented as median with IQR and, normally distributed data are presented in box plots and error bars represent mean with SD. For statistical analysis Kruskall Wallis or one-way ANOVA (when appropriate) was performed and significant differences were indicated as *p<0.05 or **p<0.01. All symbols represent data from single individuals (• healthy controls (HC), ^▄^ RA-risk individuals (RA at-risk), Δ early RA patients (RA).

Since IL-21 and IL-10 production was below detection limit in unstimulated cells, we analysed IL-21 and IL-10 production in LN CD4^+^CXCR5^+^ and CD8^+^CXCR5^+^ Tfh cells upon ex vivo stimulation with PMA/ionomycin. Among CD4^+^CXCR5^+^ Tfh cells, the frequency of IL-21 producing cells upon ex vivo stimulation was significantly decreased in early RA patients (p<0.01) when compared with HCs, while the frequency was intermediate for the RA-risk individuals (Figure 2D). Similar findings were observed for CD8^+^CXCR5^+^ Tfh cells, but without reaching statistical significance (Figure 2D). In contrast, the frequency of IL-10 producers in the CD4^+^ and CD8^+^ CXCR5^+^ Tfh cells was comparable between the three study groups (Figure 2E).

### In lymphoid tissue, the frequencies of CD4^+^ and CD8^+^ follicular helper T cells correlate with the frequencies of CD19^+^ B cells

Since CD4^+^CXCR5^+^ and CD8^+^CXCR5^+^ Tfh cells were increased in LNs of RA patients compared to HCs, we investigated if they were in any way related to the frequencies of B cells found in LNs. Consistent with previous findings (29), the frequency of CD19^+^ B cells was increased in early RA patients compared with healthy controls (p<0.05) but similar between healthy controls and RA-risk individuals (Figure 3A). When compared with CD4^+^CXCR5^+^ Tfh cells, we found a strong and significant correlation between the frequencies of CD19^+^ B cells and CD4^+^CXCR5^+^ Tfh cells (p<0.0001, r=0.76) (Figure 3B). Interestingly, we also observed a significant and strong correlation (p<0.0004, r=0.62) between CD8^+^CXCR5^+^ Tfh cells with CD19^+^ B cells in LN (Figure 3C). Taken together, the frequencies of CD4^+^CXCR5^+^ and CD8^+^CXCR5^+^ Tfh cells strongly correlate with the frequencies of CD19^+^ B cells localised within LNs.

**Figure 3.**
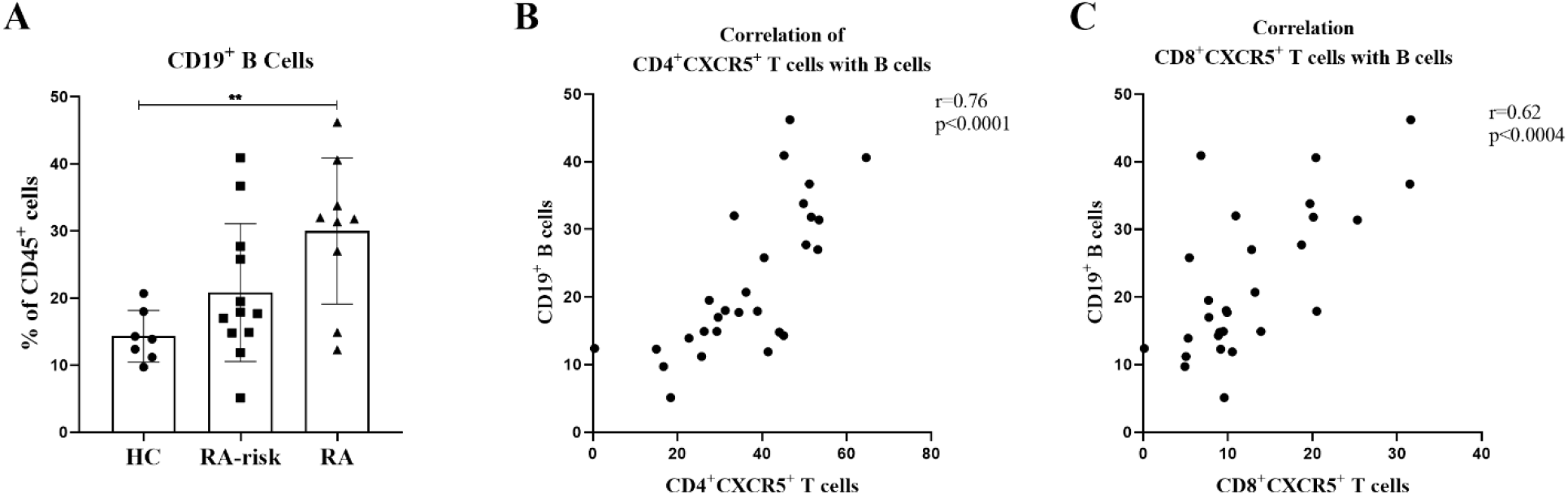
Correlation of CD19^+^ B cells with CD4^+^ and CD8^+^ follicular helper T cells in LNs. **(A)** frequencies of CD19^+^ B cells in LNs is shown for HCs, RA-risk and early RA patients. **(B)** Correlation between the frequencies of CD4^+^CXCR5^+^ Tfh cells and the frequencies of CD19^+^ B cells in LN. **(C)** Correlation between the frequencies of CD8^+^CXCR5^+^ Tfh cells and the frequencies of CD19^+^ B cells in LNs. Correlations are shown for the total group. Error bars represent mean with SD. For statistical analysis Kruskall Wallis or one-way ANOVA (when appropriate) was performed and significant differences were indicated as *p<0.05 or **p<0.01. All symbols represent data from single individuals (• healthy controls (HC), ^▄^ RA-risk individuals (RA at-risk), Δ early RA patients (RA).

## DISCUSSION

We studied CD4^+^ and CD8^+^ Tfh cells in both LN biopsies and peripheral blood samples obtained during the earliest phases of RA and compared our findings to control samples. We found an increased frequency of CD4^+^CXCR5^+^ follicular helper T cells and CD19^+^ B cells in lymphoid tissue of early RA patients, and CD8^+^CXCR5^+^ Tfh cells was similarly increased in LN tissue of both RA-risk individuals and RA patients. This increased frequency of B cells in LNs of RA patients compared to healthy controls is in accordance with previous reports (29, 33). A plausible explanation behind this increase could be the retention of B cells in lymphoid tissue where they eventually differentiate into various B cell effector phenotypes and contribute to disease development. In LN biopsies the frequencies of CD4^+^CXCR5^+^ and CD8^+^CXCR5^+^ Tfh cells correlate significantly with the frequency of CD19^+^ B cells suggesting an increased number of B and T cells that can possibly interact at the LN follicular border and drive immune responses. In mice, the location of CD4^+^ Tfh cells inside the LN during primary and memory responses has been studied in detail (34). During the primary immune response CD4^+^ Tfh cells are located in the GC follicle, while for a memory response CD4^+^ Tfh cells are located in the sub capsular region and can leave the follicle via the lymphatic flow. This enables CD4^+^ Tfh cells to migrate through blood towards other secondary lymphoid organs or inflamed tissues where they can initiate new GC responses if the antigen is present (14). These migrating CD4^+^ Tfh cells in blood can be present before differentiation towards a fully mature effector Tfh phenotype (35). In our study, the frequency of active and quiescent CD4^+^ Tfh cells in blood was not significantly altered in RA and RA-risk individuals compared with HCs.

CD8^+^ T cells often function as an effector cytotoxic T cell type hence, they were assumed to be excluded from entry into B cell follicles and participation in GC reactions (36, 37). However, recent data suggest that CD8^+^ T cells like their CD4^+^ counterpart are able to acquire CXCR5 which enables their migration into B cell follicles and subsequently eliminate virus infected B cells as well as CD4^+^ Tfh cells (19). Our findings on the presence and increased frequency of CD8^+^ Tfh cells in lymphoid tissues of early RA patients confirm the presence of CD8^+^ Tfh cells in lymphoid structures. A study by Kang *et al* was among the first to report the presence of CD8^+^ T cells in ectopic lymphoid follicles in joints of RA patients (38). In addition, a recent study reported an increased frequency of CD8^+^CXCR5^+^PD1^+^ Tfh cells in LN tissues of IL-2 knock-out mice which were shown to secrete IL-21 and promote B cell antibody class-switching. Interestingly, these CD8^+^ T follicular cells continued to expand in terms of frequency and numbers over time in these mice (20). Although we did not investigate the B cell class-switching potential of the observed lymph node CD8^+^ Tfh cells, their frequency correlated with the numbers of B cells. Human studies are needed to further unravel the potential of these cells to participate in a typical GC reaction such as B cell selection.

Even though Tfh cells are thought to be crucial for B differentiation within GCs, a recent study identified another type of B-cell promoting T helper cell in peripheral blood and synovial tissue expressing high levels of PD-1 while negative for CXCR5 (26). These cells, identified as peripheral helper cells (Tph), express factors like IL-21, CXCL13 and ICOS enabling them to provide B cell help and drive B cell differentiation. Further characterisation of Tph and Tfh cells in lymphoid tissues as well as inflamed synovial tissue is of interest to understand the mechanisms leading to the formation of secondary and tertiary lymphoid structures and subsequent autoimmune responses.

Similar to previous findings of decreased production of pro-inflammatory cytokines upon ex vivo stimulation of LN T cells of early RA patients (39, 40), we found decreased production of IL-21 in LN Tfh cells of early RA patients and in part of the RA-risk individuals. This decreased production of IL-21 by CD4^+^CXCR5^+^ and CD8^+^CXCR5^+^ Tfh cells could be a consequence of prolonged antigen stimulation and exposure *in vivo*, resulting in an exhausted phenotype of these cells (41). Interestingly and in accordance with previous reports (39), this phenomenon of decreased IL-21 production by T cells in our study was only observed in LNs but not in peripheral blood. Our results further highlight the possibility that LNs may offer a unique environment that influences the function and phenotype of T cells resulting in disease development. Therefore, future work aimed at unraveling the possibility of follicular helper T cell exhaustion in lymphoid structures is needed. While IL-21 is described as the most important cytokine derived from Tfh cells to promote B-cell differentiation and proliferation, Tfh cells also produce other cytokines like IL-4 and IFN-y. Indeed, an in vitro study using sorted CD4^+^CXCR5^+^ blood Tfh cells from healthy controls (HCs) and chronic hepatitis C infected patients (HCV) showed that Tfh cells from HCV patients produce less IL-21 but are equally capable of driving *in vitro* B cell proliferation and differentiation into antibody producing cells compared with healthy control cells (42). This indicates that B-cell differentiation is not solely dependent on large amounts of IL-21. Another explanation for low ex vivo IL-21 production may be that primary immune responses are depending on IL-21 but that secondary memory responses of Tfh cells are not (42, 43).

Even though the number of participants in our study is limited, the data presented here show for the first time the relation between the number of Tfh cells and B cells inside human LN biopsies and increased frequencies of CD4^+^ and CD8^+^ Tfh cells in LN biopsies of early RA patients. Since Tfh may amplify B-cell responses and autoantibody production in lymphoid tissue and CD8^+^ Tfh are already increased in RA-risk individuals, targeting Tfh cells early could be tested as a new approach to prevent further disease progression during the earliest phases of RA.

## Acknowledgements

We thank the study participants in the study, the radiology department at the Academic Medical Center (AMC) for lymph node sampling; the flow cytometry facility at the Hematology department at AMC, especially J.A. Dobber; and the core facility Cellular Imaging of the Amsterdam UMC, location AMC.

## Author contributions

Study conception and design: LGB, DMG, PPT. Acquisition of data: THR, JH, BK, NS, KPL, MM. Analysis and interpretation of data: THR, DCA, PPT, DMG, NdV, LGB. All authors revised the article for important intellectual content and all authors have read and approved the final version of the manuscript. Dr. van Baarsen had full access to all of the data in the study and takes responsibility for the integrity of the data and the accuracy of the data analysis.

## FIGURES

**Supplementary Figure 1.**
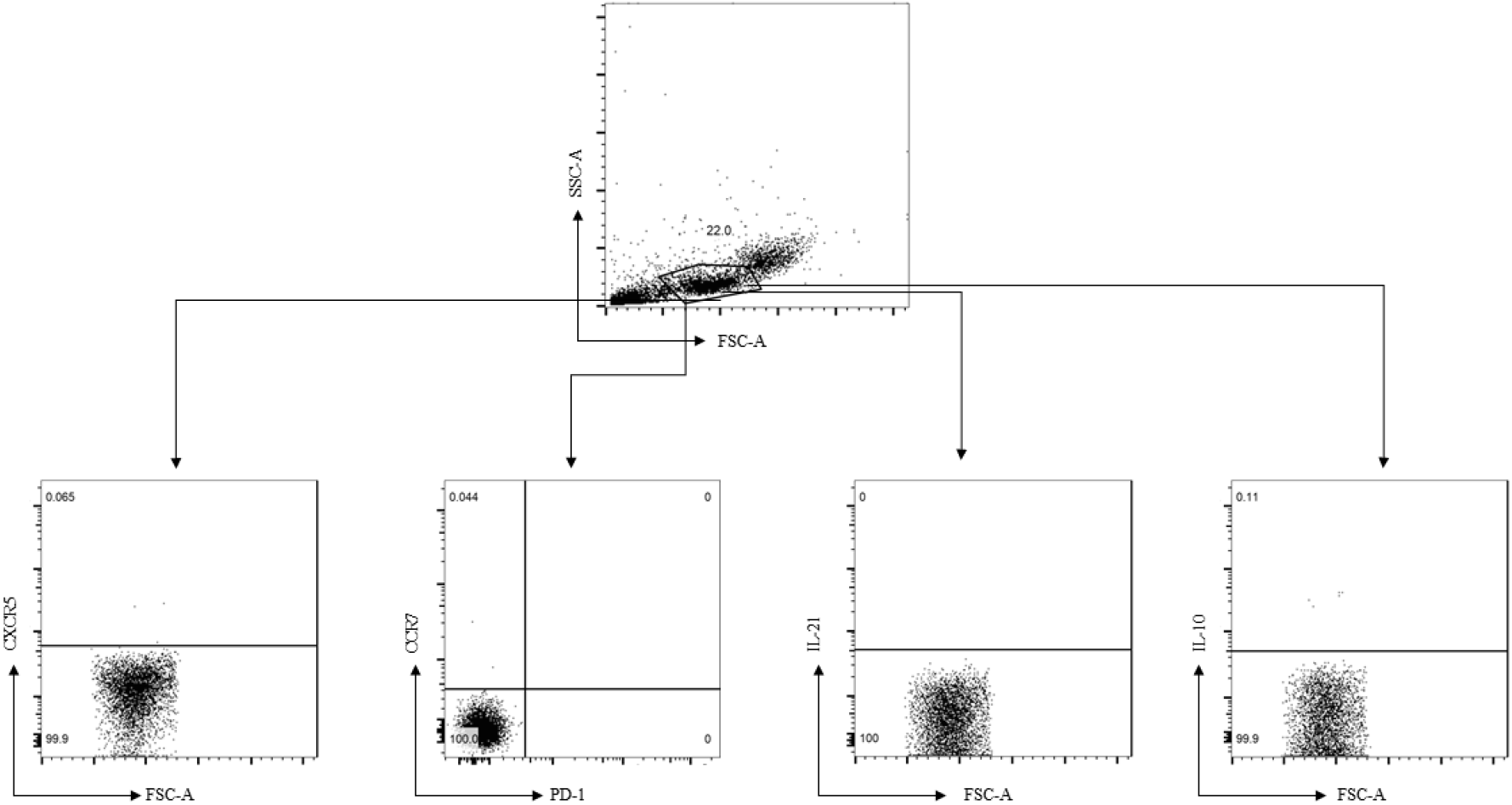
Negative control (unstained sample) used for setting of gates for the analysis of CXCR5, CCR7, PD-1, IL-21 and IL-10 cells in PBMCs.

